# Interstitial Notch signaling regulates nephron development via the Gata3-Renin axis in the mouse kidney

**DOI:** 10.1101/2022.06.08.495282

**Authors:** Eunah Chung, Mike Adam, Andrew S. Potter, Sara M. Marshall, S. Steven Potter, Joo-Seop Park

## Abstract

Notch signaling in the renal interstitium is known to be required for the formation of mesangial cells and *Ren1* (Renin)-expressing cells. However, little is known about how interstitial Notch signaling affects nephron development. We found that blocking Notch signaling in the renal interstitium in mice caused developmental arrest of proximal tubules accompanied by defective formation of mesangial cells. We examined the interstitial *Pdgfrb* mutant kidney which exhibits a similar mesangial cell defect and found that the *Pdgfrb* mutant kidney showed normal proximal tubule development, suggesting that the absence of mesangial cells was not the cause of defective proximal tubule development. Our single cell RNA-seq analysis of the interstitial *Rbpj* mutant kidney showed that a subset of proximal tubule genes were downregulated in the mutant kidney and that *Gata3* was downregulated in the mutant interstitium during the development of *Ren1*-expressing cells. We found that deleting *Gata3* in the interstitium caused the loss of Renin and the developmental arrest of proximal tubules, phenocopying the interstitial *Notch/Rbpj* mutants. Our results suggest that interstitial Notch signaling regulates the development of proximal tubules via the Gata3-Renin axis in the mouse kidney.

## INTRODUCTION

Mesenchymal nephron progenitors expressing the transcription factor Six2 are multipotent (1). Upon the downregulation of Six2, they undergo mesenchymal-to-epithelial transition (MET) and form a renal vesicle, which eventually develops into the nephron with distinct segments (2). It is well established that both self-renewal and commitment of Six2+ cells require Wnt9b signal from the adjacent collecting duct (3-6). In addition, it was recently shown that balancing self-renewal and commitment of Six2+ cells requires cortical interstitial cells (marked by Foxd1) surrounding Six2+ cells (7). When cortical interstitial cells and their descendants were removed by *Foxd1Cre*-mediated activation of diphtheria toxin A, MET of Six2+ cells was inhibited and Six2+ cell population was expanded. While this result suggests that interstitial cells provide signals for early commitment of Six2+ cells, little is known about how these interstitial cells contribute to the development of epithelial nephron progenitors into the mature nephron.

Notch signaling, one of the major pathways mediating cell fate decisions during development, requires cell-cell contact for its activation because both the receptor and ligand are bound to the plasma membrane (8). Upon activation, the intracellular domain of a Notch receptor (NICD) is released from the membrane, enters the nucleus, forms a complex with Rbpj, and regulates the expression of target genes (8). Notch signaling plays critical roles in multiple lineages in the developing kidney. In the nephron lineage, Notch signaling is required for the timely downregulation of Six2 and the formation of all nephron segments (9, 10). In the collecting duct lineage, it regulates the binary cell fate decision between principal cells and intercalated cells (11, 12). In the interstitial lineage, Notch signaling is required for the formation of mesangial cells and juxtaglomerular cells (13-15). However, it is unknown how Notch signaling in one lineage affects the developmental processes in another lineage.

Here we show that interstitial Notch signaling is required for the development of the nephron. Conditional deletion of *Notch* or *Rbpj* by *Foxd1Cre* causes the developmental arrest of proximal tubules accompanied by defective formation of mesangial cells. Conditional deletion of *Pdgfrb* by *Foxd1Cre* causes defective formation of mesangial cells without inhibiting proximal tubule development, suggesting that the mesangial cell defect is not an underlying cause for the proximal tubule defect. Our single cell RNA-seq (scRNA-seq) analysis shows that interstitial Notch signaling is required for the expression of *Gata3* during the formation of juxtaglomerular cells. *Foxd1Cre*-mediated deletion of *Gata3* causes not only the absence of *Ren1* (encoding Renin)-expressing juxtaglomerular cells but also the developmental arrest of proximal tubules. These results suggest that the absence of the Gata3-Renin axis is responsible for the proximal tubule defect seen in the interstitial *Notch/Rbpj* mutant kidneys. Our findings are consistent with the idea that low blood flow causes the absence or paucity of mature proximal tubules as seen in a human disease known as renal tubular dysgenesis (16, 17).

## RESULTS

### Notch1 and Notch2 act redundantly in the formation of mesangial cells

It has been reported that *Foxd1Cre*-mediated deletion of *Rbpj*, a gene encoding the only DNA binding partner for Notch, in the interstitial lineage blocked the formation of mesangial cells (13). The same report showed, however, that the deletion of either *Notch1* or *Notch2* by *Foxd1Cre* did not cause the mesangial cell defect (13). To test if Notch1 and Notch2 act redundantly in the formation of mesangial cells, we deleted both *Notch1* and *Notch2* in the interstitial lineage with *Foxd1Cre*. In Figure 1A, we show that the *Notch1*/*Notch2* double mutant kidney recapitulated the mesangial cell defect seen in the interstitial *Rbpj* mutant kidneys as reported previously (13, 14). To quantitate the mesangial cell defect, we scored glomeruli into three categories: normal glomeruli (Class I), glomeruli with few or no mesangial cells (Class II), and cystic glomeruli without mesangial cells (Class III). We found that the presence of one *Notch2* allele (*Notch1*^*c/c*^*;Notch2*^*c/+*^*;Foxd1*^*GC/+*^) was sufficient for the formation of mesangial cells without any noticeable defect and that the presence of one *Notch1* allele (*Notch1*^*c/+*^*;Notch2*^*c/c*^*;Foxd1*^*GC/+*^) caused 70% of glomeruli to belong to Class II (Figure 1B). We observed the Class III defect only in the *Notch1*/*Notch2* double mutant kidney (Figure 1B). These data suggest that Notch1 and Notch2 act redundantly in the formation of mesangial cells and that Notch2 is the major Notch receptor in this process.

**Figure 1.**
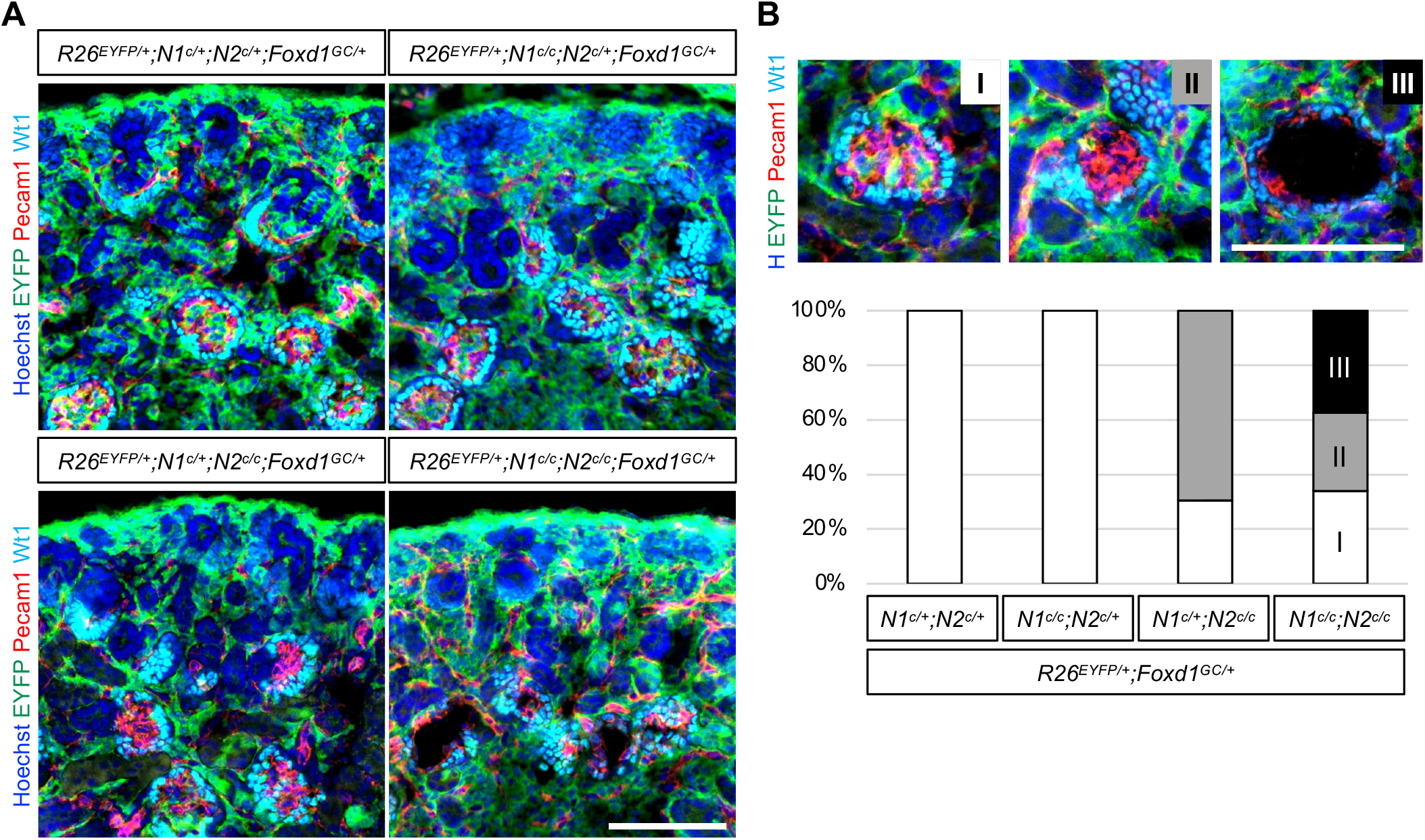
*Notch1* and *Notch2* act redundantly in the formation of mesangial cells. Stage E18.5; Scale bar 100 µm (A) *Foxd1GFPcre* (*Foxd1*^*GC*^) deletes *Notch1* and *Notch2* and activates *Rosa26-EYFP* reporter in the interstitium. Deletion of *Notch1* causes no defect in mesangial cells. Deletion of *Notch2* causes the absence of EYFP^+^ interstitial cells in a subset of glomeruli. The mesangial cell defect is the most severe in the *Notch1*/*Notch2* double mutant kidney. (B) Quantitative scoring of the mesangial cell defect. Class I represents normal glomeruli containing both Pecam1^+^ endothelial cells (red) and EYFP^+^ mesangial cells (green). Class II represents glomeruli containing endothelial cells with a minimal presence of mesangial cells. Class III represents cystic glomeruli without mesangial cells, which can be seen in the *Notch1*/*Notch2* double mutant kidney only. More than 100 glomeruli were scored for each genotype. Results are representative of three independent experiments.

### Interstitial Notch signaling is required for the development of proximal tubules

Unexpectedly, we found that proximal tubule development was defective in the *Notch1*/*Notch2* double mutant kidney. Fewer *Lotus tetragonolobus* lectin (LTL)-stained cells were present and the LTL signal was weaker in the double mutant kidney (Figure 2A). It has been established that staining for Cdh6 and LTL enables us to stage proximal tubule development, owing to the fact that Cdh6 and LTL mark proximal tubule progenitors and mature proximal tubules, respectively (18-20). Immature proximal tubule progenitors show strong Cdh6 signal (Cdh6^high^) with low or no LTL signal (LTL^low^ or LTL^neg^). Mature proximal tubules show weak Cdh6 signal and strong LTL signal (Cdh6^low^ LTL^high^) (18, 19). All immature and mature proximal tubules express Hnf4a, the transcription factor required for the development of immature proximal tubules into mature proximal tubules (18, 19, 21). Only in the *Notch1*/*Notch2* double mutant kidney were mature proximal tubules absent, indicating that Cdh6^high^ LTL^low^ cells failed to develop into Cdh6^low^ LTL^high^ cells (Figure 2B). To further examine the role of interstitial Notch signaling on the development of proximal tubules, we have generated the interstitial *Rbpj* mutant kidney with *Foxd1Cre*. We found that, when interstitial Notch signaling was blocked by the loss of Rbpj, immature proximal tubule cells failed to develop into mature proximal tubules (Figure 3), phenocopying the Notch1/2 double mutant kidney (Figure 2). These data strongly suggest that the formation of mature proximal tubules require interstitial Notch signaling.

### Loss of Rbpj in the interstitial lineage results in the downregulation of Pdgfrb in the interstitium and Wt1 in podocytes

The mesangial cell defect seen in the *Rbpj* or *Notch1*/*Notch2* double mutant kidneys by *Foxd1Cre* was reminiscent of the phenotype of the *Pdgfrb* mutant kidney as previously described (22), raising the possibility that Notch signaling may regulate the expression of *Pdgfrb* in the interstitium. We found that, in the control kidney, *Pdgfrb* is widely expressed in the renal interstitium. Its expression was particularly high in mesangial cells and interstitial cells adjacent to nascent nephron structures and thick blood vessels (Figure 4). In the *Rbpj* mutant kidney, *Pdgfrb* was still expressed but fewer Pdgfrb^high^ cells were present. Pdgfrb signal in the mutant glomeruli was noticeably weaker or absent, consistent with the notion that interstitial Notch signaling is required for the formation of mesangial cells (13, 14). These results suggest that interstitial Notch signaling is dispensable for the expression of *Pdgfrb* but required for the upregulation of *Pdgfrb* in interstitial cells adjacent to endothelial cells. We found that, in the *Rbpj* mutant kidney, Wt1 signal in the nascent glomeruli closer to the nephrogenic zone appeared normal but the more developed glomeruli located farther away from the nephrogenic zone (marked by while arrowheads in Figure 4) showed weaker Wt1 signal. This finding suggests that the presence of mesangial cells is required for the maintenance of *Wt1* expression in podocytes.

**Figure 2.**
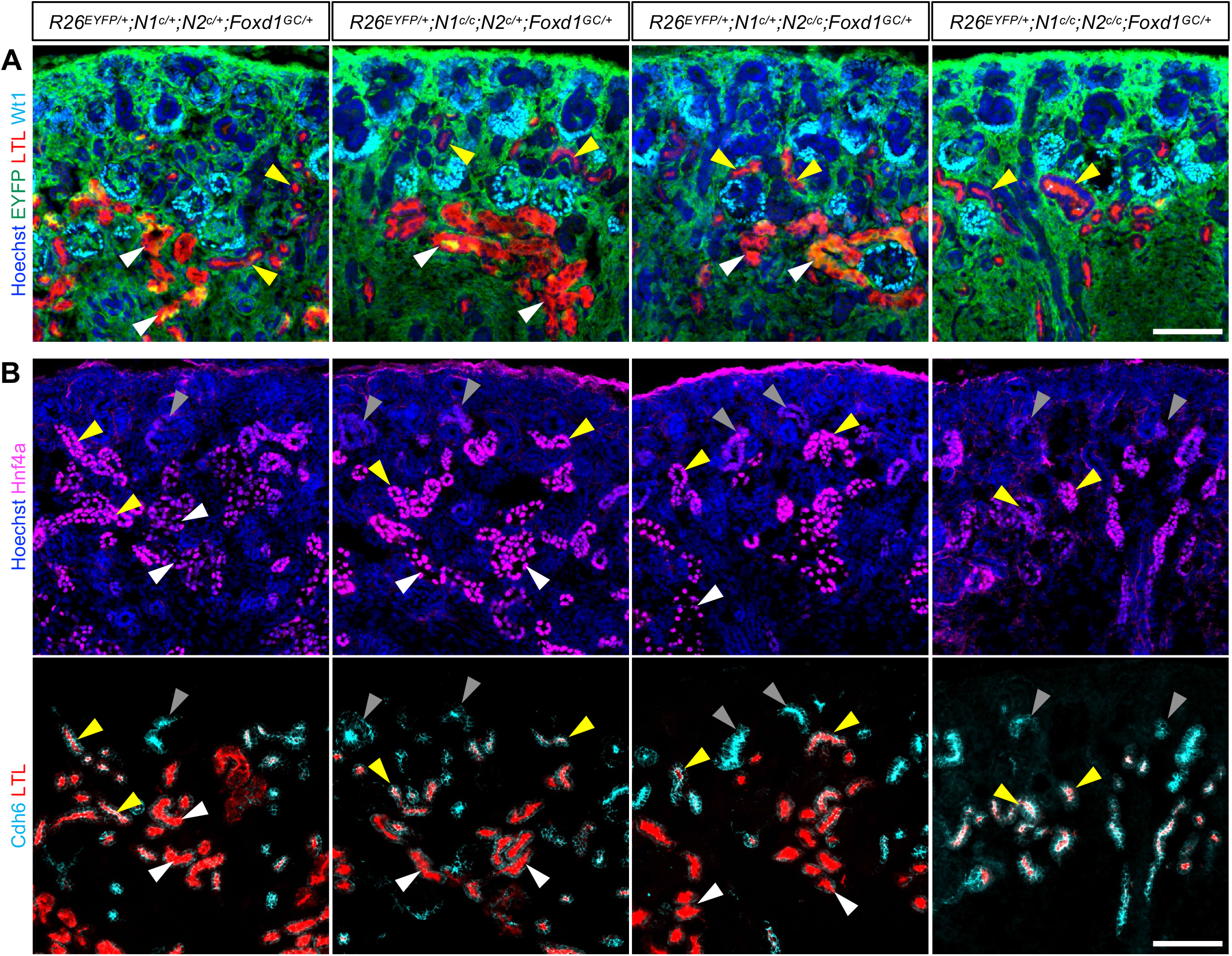
Interstitial Notch signaling is required for the development of proximal tubules. Stage E18.5; Scale bar 100 µm (A) *Wt1* expression is strong in developing podocytes and weak in the cap mesenchyme. Immature proximal tubules (marked by yellow arrowheads) show LTL staining in only the luminal side of the epithelia. Mature proximal tubules (marked by white arrowheads) show strong LTL staining throughout the entire cell body. In the *Notch1*/*Notch2* double mutant kidney, only luminal LTL staining is detected. (B) Hnf4a is detected in all developing proximal tubules. Gray arrowheads mark Cdh6^high^ LTL^neg^ cells and yellow arrowheads mark Cdh6^high^ LTL^low^ cells. The most mature proximal tubules (Cdh6^low^ LTL^high^, marked by white arrowheads) are absent in the *Notch1*/*Notch2* double mutant kidney. Images are representative of three independent experiments.

**Figure 3.**
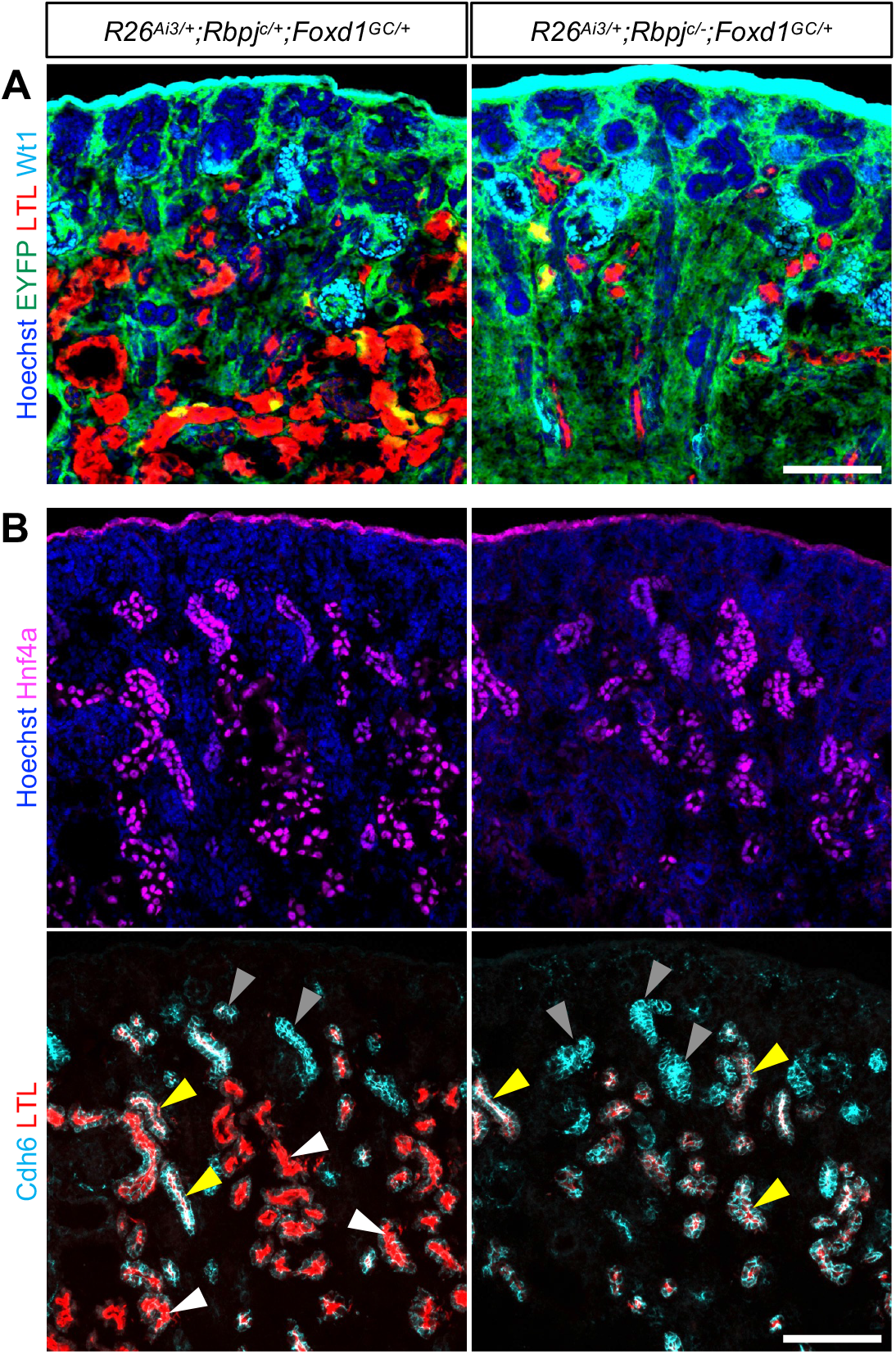
Loss of Rbpj in interstitial cells causes the developmental arrest of proximal tubules. Stage E18.5; Scale bar 100 µm (A) LTL staining is weaker in the *Rbpj* mutant kidney. (B) The most mature proximal tubules (Cdh6^low^ LTL^high^, marked by white arrowheads) are absent in the *Rbpj* mutant kidney. Gray arrowheads mark Cdh6^high^ LTL^neg^ cells and yellow arrowheads mark Cdh6^high^ LTL^low^ cells. Images are representative of four independent experiments.

**Figure 4.**
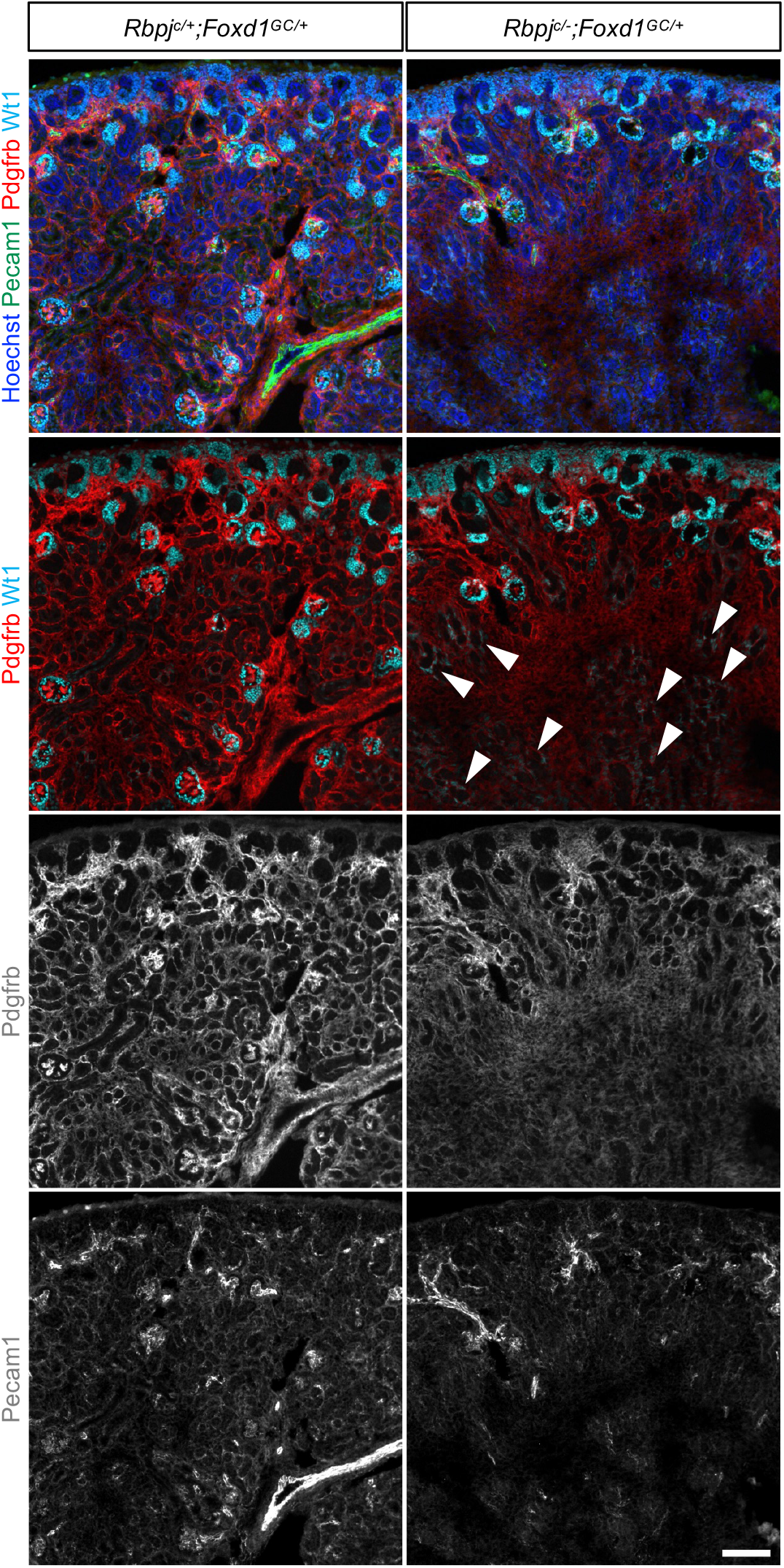
Loss of Rbpj in the interstitial lineage results in the downregulation of *Pdgfrb* in the interstitium and *Wt1* in podocytes. In the control kidney, Pdgfrb^high^ cells are adjacent to endothelial cells in the nephrogenic zone and glomeruli. In the mutant kidney, fewer Pdgfrb^high^ cells are present and glomeruli located deeper in the kidney (marked by white arrowheads) show weaker Wt1 signal. Stage E18.5; Scale bar 100 µm. Images are representative of two independent experiments.

### Loss of Pdgfrb in interstitial cells does not cause the developmental arrest of proximal tubules

In order to investigate if the absence of mesangial cells has an effect on the development of proximal tubules, we examined another mutant kidney with mesangial cell defects. It has been shown that mesangial cells are absent in the *Pdgfrb* mutant kidney (22) and we reasoned that, if the absence of mesangial cells is a cause for the proximal tubule defect, the *Pdgfrb* mutant kidney should exhibit the same defect. We generated the *Pdgfrb* mutant kidney by *Foxd1Cre* and found that *Foxd1Cre* removed most of the Pdgfrb signal from the interstitium but Pdgfrb^+^ cells were still present in a subset of glomeruli (Figure 5A, yellow arrowheads). This was likely due to mosaic action of *Foxd1Cre*, namely some interstitial cells escaped the *Foxd1Cre*-mediated deletion of *Pdgfrb* (Figure 5A) and still carried the intact *Pdgfrb* allele. We saw that, despite the absence of mesangial cells (Figure 5B, white arrowheads), mature proximal tubules (Cdh6^low^ LTL^high^) were formed in the *Pdgfrb* mutant kidney (Figure 5C). These results suggest that the absence of mesangial cells is not a direct cause for defective proximal tubule development.

**Figure 5.**
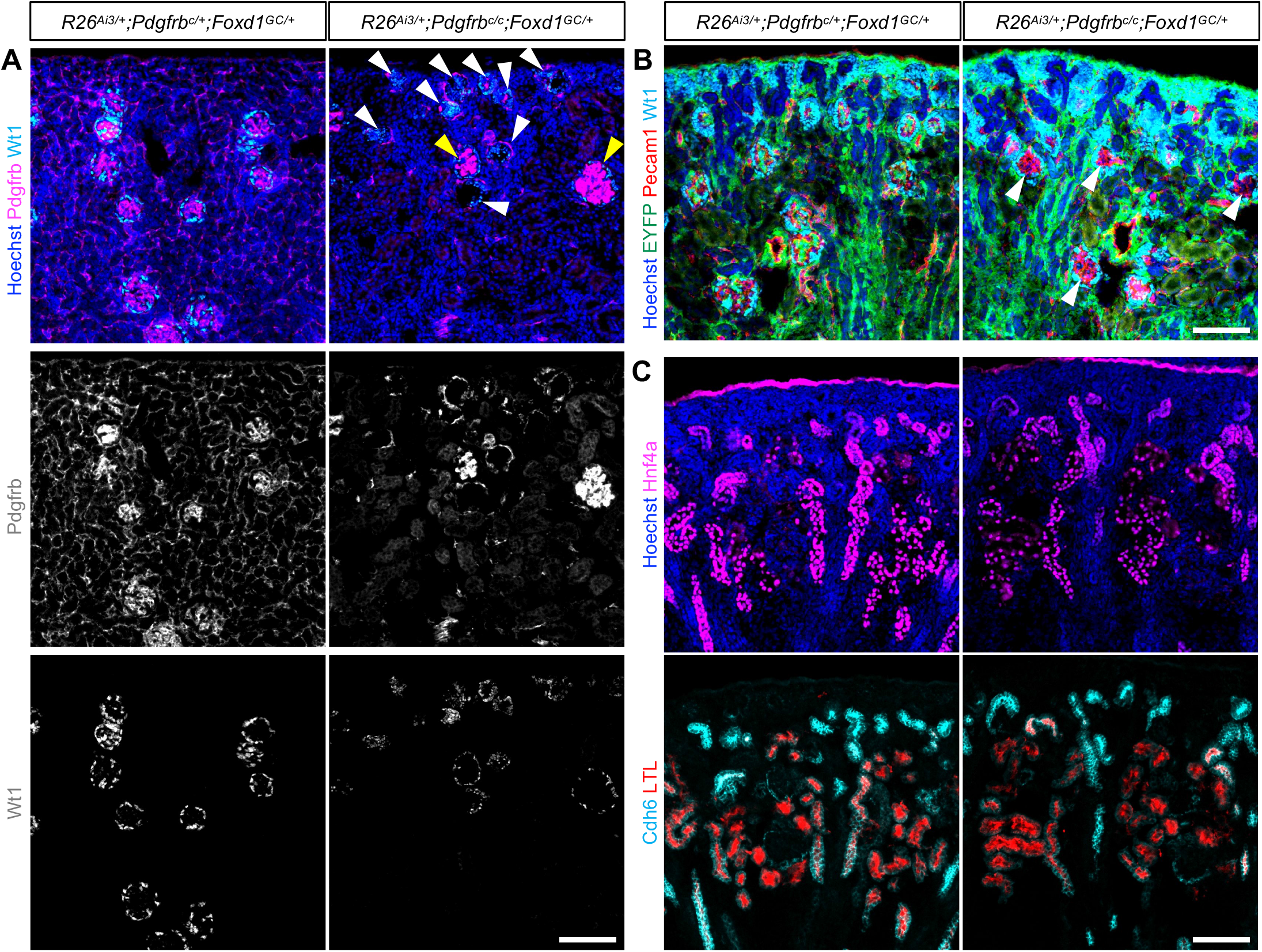
Loss of Pdgfrb in interstitial cells does not cause the developmental arrest of proximal tubules. Scale bar 100 µm (A) In the mutant kidney, Pdgfrb^+^ cells are still present, suggesting that *Foxd1Cre* is mosaic. While a subset of glomeruli have strong Pdgfrb signal (marked by yellow arrowheads), most of the glomeruli appear small and defective (marked by white arrowheads). Stage P15. (B) The *Pdgfrb* mutant kidney shows the mesangial cell defect (marked by white arrowheads). Stage P0. (C) Proximal tubule development appears normal in the *Pdgfrb* mutant kidney. Stage P0. Images are representative of two independent experiments.

### scRNA-seq analysis of the interstitial *Rbpj* mutant kidney

To further investigate how the loss of interstitial Notch signaling affects nephron and interstitial lineages in the developing kidney, we performed single cell RNA-seq (scRNA-seq) analysis of the interstitial *Rbpj* mutant and control kidneys at embryonic day 18 (E18.5). (Supplemental Figure 1 & Supplemental Table 2). Consistent with the developmental arrest of proximal tubules shown in Figure 3, we found that a subset of proximal tubule marker genes were downregulated in the mutant kidney (Figure 6A). *Heyl*, known to be a Notch target gene in many cell types (8), was found to be expressed in multiple domains in the control interstitium and downregulated in the mutant interstitium (Figure 6B, circled in green), confirming that interstitial Notch signaling was indeed blocked by the loss of Rbpj. Consistent with previous reports that Notch signaling is required for the expression of *Ren1* (14) and that Renin^+^ cells express *Gata3* (23), our scRNA-seq data showed that *Ren1* and *Gata3* were expressed in juxtaglomerular cells in the control kidney and that both of those genes were downregulated in the mutant kidney (Figure 6B, circled in blue). These results raise the interesting possibility that Notch signaling may regulate the expression of *Ren1* through Gata3.

**Figure 6.**
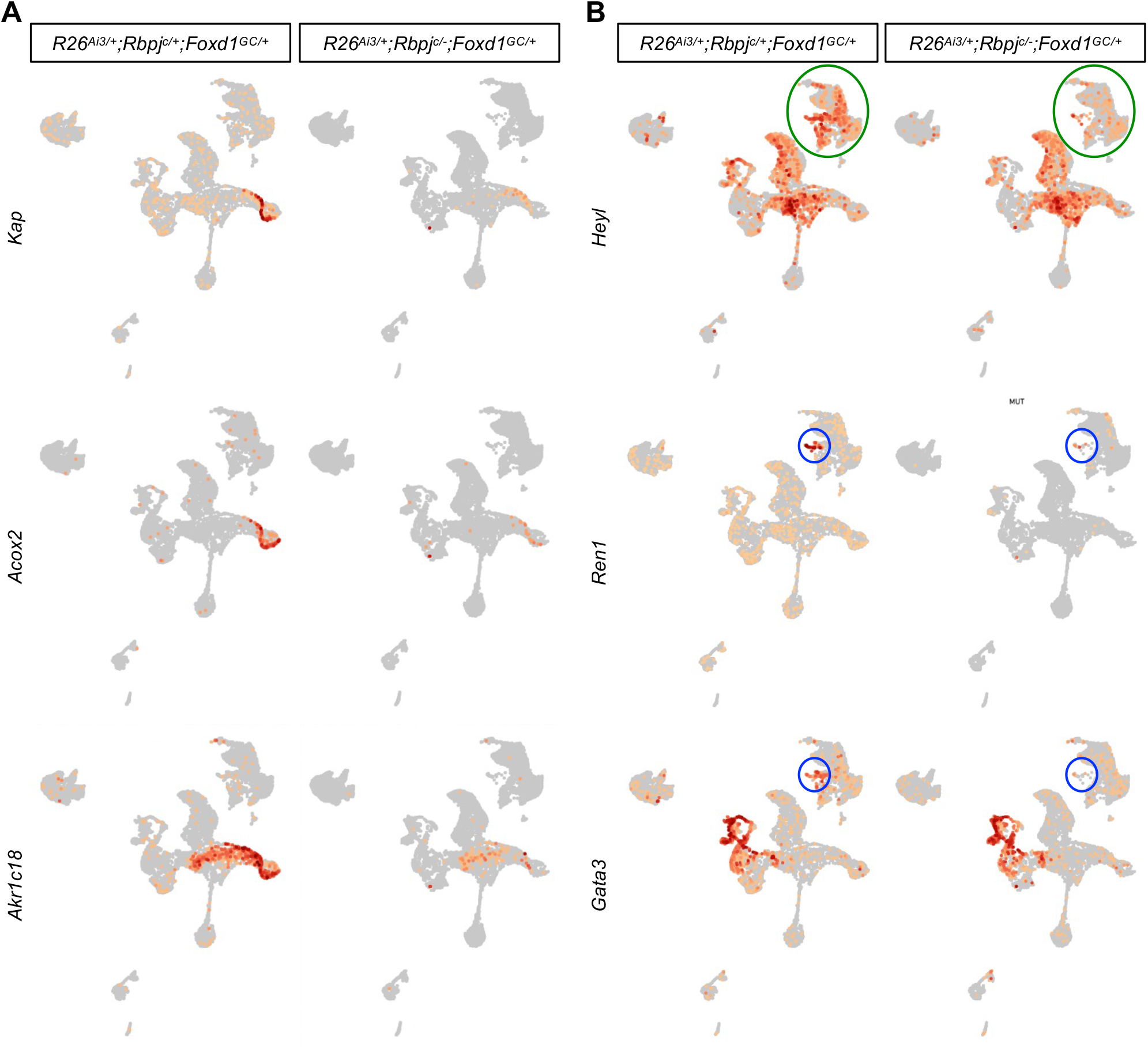
scRNA-seq analysis of the interstitial *Rbpj* mutant and control kidneys at E18.5 (A) A subset of proximal tubule marker genes are downregulated in the interstitial *Rbpj* mutant kidney (B) *Heyl*, a classic Notch target gene, is downregulated in the mutant interstitium (circled in green). *Ren1* and *Gata3* are expressed in juxtaglomerular cells (circled in blue) in the control kidney and they are downregulated in the mutant kidneys.

### Loss of Gata3 in interstitial cells causes the ablation of *Ren1* expression and the developmental arrest of proximal tubules

To test if Gata3 is required for *Ren1* expression, we conditionally deleted *Gata3* from the interstitium with *Foxd1Cre*. We found that Renin signal was ablated in the *Gata3* mutant kidney, suggesting that Gata3 is required for the expression of *Ren1* (Figure 7A). Next, we examined the development of proximal tubules and found that mature proximal tubules were absent in the *Gata3* mutant kidney (Figure 7B). Given that severe low blood pressure caused by defective Renin-Angiotensin system results in the absence or paucity of mature proximal tubules in renal tubular dysgenesis (OMIM 267430) (16, 17), our data suggest that Gata3-mediated expression of *Ren1* is required for the development of proximal tubules and that the absence of the Gata3-Renin axis is the underlying cause for the proximal tubule defect seen in the interstitial *Notch/Rbpj* mutant kidneys.

**Figure 7.**
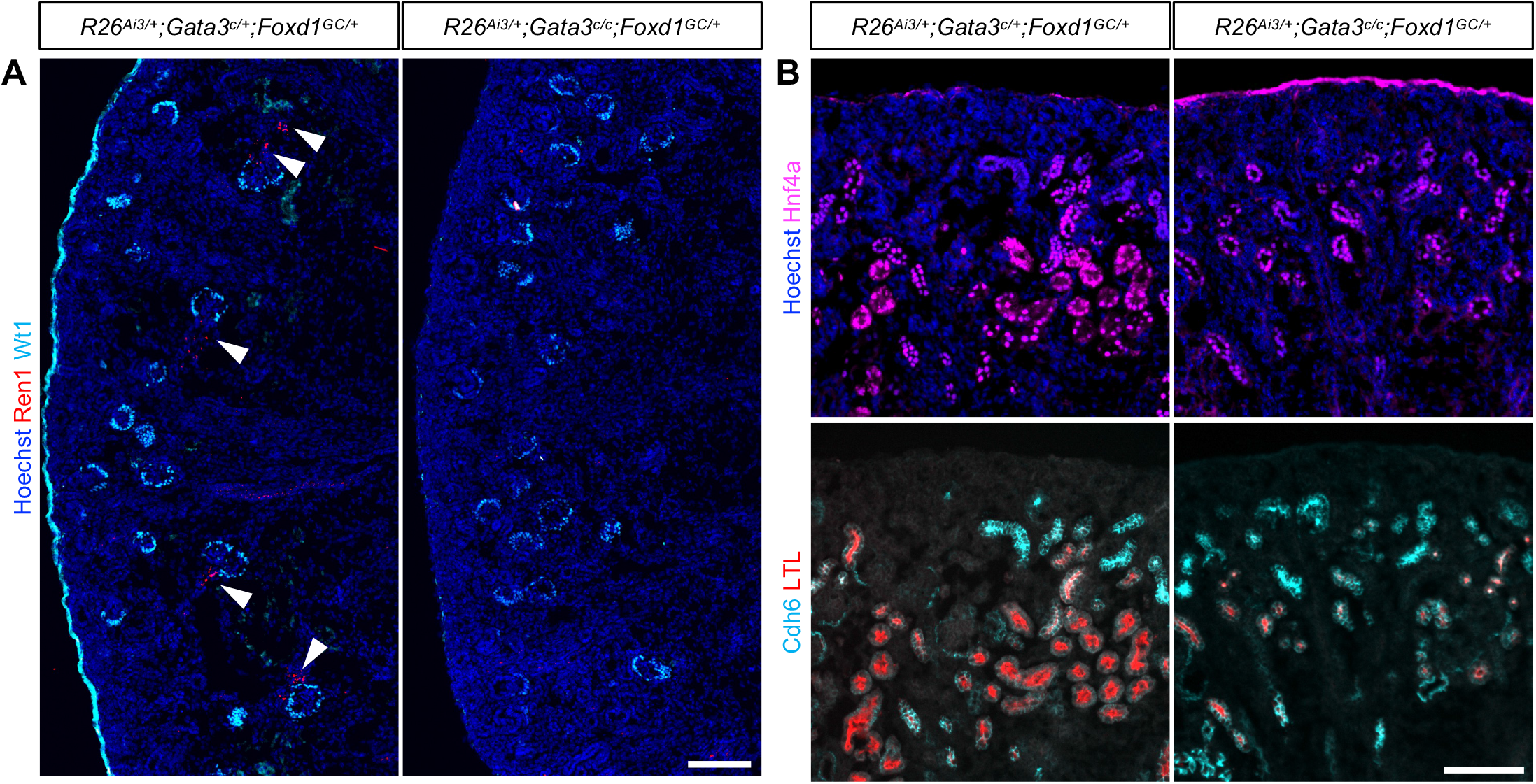
Loss of Gata3 in interstitial cells causes the ablation of *Ren1* expression and the developmental arrest of proximal tubules. Stage P0; Scale bar 100 µm (A) In the control kidney, Renin is detected in juxtaglomerular cells (marked by white arrowheads). In the *Gata3* mutant kidney, Renin signal is absent, suggesting that interstitial expression of *Gata3* is required for *Ren1* expression. (B) In the *Gata3* mutant kidney, mature proximal tubules (Cdh6^low^ LTL^high^) are absent, suggesting that the interstitial expression of *Gata3* is required for the development of proximal tubules. Images are representative of four independent experiments.

## DISCUSSION

Renin-expressing cells are known as juxtaglomerular cells because they are located adjacent to the afferent arteriole entering the glomerulus (2). They play critical roles in maintaining blood pressure. Defective Renin-Angiotensin function in humans causes renal tubular dysgenesis whose histopathologic hallmark is the absence of mature proximal tubules (16, 17). Remarkably, it is not understood how a defective Renin-Angiotensin axis would affect proximal tubule development in mouse models.

Several lines of evidence suggest that the absence of mesangial cells seen in the interstitial *Notch/Rbpj* mutant kidneys is not the underlying cause for the absence of mature proximal tubules: (1) The interstitial *Pdgfrb* mutant kidneys, despite their mesangial cell defect, showed normal proximal tubule development, (2) Deletion of two Notch2 alleles and one Notch1 allele from the interstitium (*Notch1*^*c/+*^*;Notch2*^*c/c*^*;Foxd1*^*GC/+*^) caused the mesangial cell defect but not the proximal tubule defect, (3) In the interstitial *Notch1/Notch2* double mutant kidney (*Notch1*^*c/c*^*;Notch2*^*c/c*^*;Foxd1*^*GC/+*^), 30% of glomeruli showed no mesangial cell defect but the proximal tubule defect appears to be uniform rather than mosaic. Taken together, these results rule out the possibility that the absence of mesangial cells leads to the absence of mature proximal tubules.

In addition to the mesangial cell defect, the interstitial *Notch/Rbpj* mutant kidneys lack Renin-expressing cells. Our data suggest that interstitial Notch signaling regulates the expression of *Ren1* via Gata3. However, an *in vitro* study suggested that Rbpj regulates *Ren1* directly (24). If the latter is the case *in vivo*, Notch/Rbpj complex may cooperate with Gata3 on the enhancer regulating the expression of *Ren1*. To address this, *in vivo* mapping of Gata3 and Notch/Rbpj in juxtaglomerular cells would be crucial. Given a previous report that the *Gata3* heterozygous kidney had fewer mesangial cells in the glomeruli (23), we were surprised that the mesangial cell defect in the interstitial *Gata3* mutant kidney was not as severe as that in the interstitial *Notch/Rbpj* or *Pdgfrb* mutant kidneys (Supplemental Figure 2). The simplest explanation for this is that Gata3 regulates the proliferation of mesangial cells rather than their migration into glomeruli.

Our mutant mice that had *Notch/Rbpj* or *Gata3* genes deleted in the interstitium failed to express *Ren1* and died shortly after birth, making it impossible to examine them postnatally. This is consistent with the fact that *Ren1* mutants also show postnatal lethality (25-27). Loss of Renin would lead to decreased glomerular filtration, resulting in reduced or absent flow inside nephron tubules. It is possible that proximal tubules, other nephron segments, and even collecting duct may be able to detect luminal flow and that their development or maturation may require mechanosensing or mechanotransduction. It is tempting to speculate that Cdh6^high^ immature proximal tubules are associated with immature glomeruli without filtration or flow while LTL^high^ mature proximal tubules are connected with mature glomeruli with active filtration. Further investigation is needed to answer this interesting question.

Our work presented here highlights the important role of Renin on the development of proximal tubules in the mouse kidney. This finding has a critical implication in kidney organoids. It is known that nephron tubules formed in kidney organoids remain immature (28, 29). Perhaps the lack of the Renin-Angiotensin system and associated circulation system is the underlying cause for the absence of mature nephron tubules in kidney organoids (30).

## METHODS

### Mice

All mouse alleles used in this study have been previously described: *Foxd1*^*GFPcre*^ (*Foxd1*^*GC*^) (31), *Rosa26*^*EYFP*^ (32), *Rosa26*^*Ai3*^ (33), *Notch1*^*c*^ (34), *Notch2*^*c*^ (35), *Rbpj*^*c*^ (36), *Rbpj*^*-*^ (37), *Pdgfrb*^*c*^ (38), and *Gata3*^*c*^ (39). All mice were maintained in the Cincinnati Children’s Hospital Medical Center (CCHMC) animal facility according to animal care regulations.

### Immunofluorescence

Kidneys were fixed in phosphate-buffered saline (PBS) containing 4% paraformaldehyde for 10-30 min, incubated at 4°C overnight in PBS containing 10% sucrose, embedded in OCT (Thermo Fisher Scientific), and stored at -80°C. Cryosections (9 μm) were incubated at 4°C overnight with primary antibodies (Supplemental Table 1) in PBS containing 5% heat-inactivated sheep serum and 0.1% Triton X-100. Fluorophore-labeled secondary antibodies were purchased from Thermo Fisher Scientific and Jackson Immuno. Images were taken with a Nikon Ti-E widefield microscope at the Confocal Imaging Core at CCHMC.

### scRNA-seq

Dissociation was performed using psychrophilic proteases (40). Briefly, tissue was incubated in digestion buffer containing 10 mg/ml *Bacillus Licheniformis* enzyme (Sigma, P5380) for 20 min, with trituration and vigorous shaking every 2 min. The digest mixture was then filtered using a 30 µM filter (Miltenyi), and the filter was rinsed with 10 ml ice-cold 10% FBS/PBS. The flow-through was centrifuged at 300 g for 5 min at 4°C. Supernatant was removed, and the pellet was resuspended in 1 ml 10% FBS/PBS, followed by filtration using a 20 µM filter (pluriSelect), rinsing the filter with 8 ml ice-cold 10% FBS/PBS, and centrifuging the flow-through at 300 g for 5 min at 4°C. The pellet was resuspended in 0.4-1 ml ice-cold 10% FBS/PBS. After the cell suspension was analyzed using a hemocytometer with trypan blue, 9600 cells were loaded into the 10X Chromium instrument for each sample (two controls and two mutants), targeting recovery of 6,000 cells, and Gel Beads in Emulsion (GEMs) were generated. 10X Genomics 3’ v3.1 chemistry was used, using the protocol provided by 10X Genomics, with 14 cycles for cDNA amplification. Single cell libraries were sequenced using the NovaSeq 6000.

### scRNA-seq data analysis

The fastq files were processed using 10x Genomics Cell Ranger v6.0.0 (10x Genomics) aligning reads to the mouse genome (mm10) to generate a feature count matrix. The four samples (two controls and two mutants) were merged into a single feature-barcode matrix using the aggr function of Cell Ranger. Cell-type clusters and marker genes were identified using the R v4.1.1 library Seurat v4.1.0 (41). Initial cell filtering selected cells that expressed >500 genes. Genes included in the analysis were expressed in a minimum of three cells. Only one read per cell was needed for a gene to be counted as expressed per cell. Cells containing high percentages of mitochondrial, >30%, and hemoglobin genes, >2.5% were filtered out. The resulting gene expression matrix was normalized using sctransform (42). The final total number of 24062 cells (control: 11530, mutant: 12532) were analyzed. All clustering was unsupervised, without driver genes. Genes with the highest variability among cells were used for principal components analysis. Cell clusters were determined by the Louvain algorithm by calculating k-nearest neighbors and constructing a shared nearest neighbor graph, with a resolution set at 0.2. Dimension reduction was performed using UMAP (Uniform Manifold Approximation and Projection) using the first twenty principal components. Marker genes were determined for each cluster using the Wilcoxon Rank Sum test within the FindAllMarkers function using genes expressed in a minimum of 25% of cells and fold change threshold of 1.3. Data are available at Gene Expression Omnibus (GEO) under accession number GSE202882.

### Study approval

All experiments were performed in accordance with animal care guidelines and the protocol was approved by the Institutional Animal Care and Use Committee of the Cincinnati Children’s Hospital Medical Center (IACUC2017-0037 and IACUC2020-0045). We adhere to the National Institutes of Health Guide for the Care and Use of Laboratory Animals.

## Author contributions

E.C. and S.M.M. performed mouse experiments. M.A., A.S.P., and S.S.P performed scRNA-seq and data analysis. E.C. and J.P. designed the experiments, analyzed the data, made figures, and cowrote the manuscript. All authors approved the final version of the manuscript.

## Acknowledgments

We thank the Confocal Imaging Core (CIC) and the DNA Sequencing and Genotyping Core (DSGC) at CCHMC. This work was supported by the National Institute of Diabetes and Digestive and Kidney Diseases, National Institutes of Health R01 DK127634, R01 DK120847, R01 DK125577, and R01 DK131052 to J.P.

## Supplemental Material

Supplemental Figure 1. Clustering of 17 cell types from scRNA-seq analysis

Supplemental Figure 2. Mesangial cell defect in the *Gata3* mutant kidney by *Foxd1Cre*

Supplemental Table 1. Antibodies used for immunofluorescence

Supplemental Table 2. Marker gene list from scRNA-seq

## REFERENCES

1. Kobayashi A, Valerius MT, Mugford JW, Carroll TJ, Self M, Oliver G, et al. Six2 defines and regulates a multipotent self-renewing nephron progenitor population throughout mammalian kidney development. Cell Stem Cell. 2008;3(2):169–81.

2. McMahon AP. Development of the Mammalian Kidney. Curr Top Dev Biol. 2016;117:31–64.

3. Carroll TJ, Park JS, Hayashi S, Majumdar A, and McMahon AP. Wnt9b plays a central role in the regulation of mesenchymal to epithelial transitions underlying organogenesis of the mammalian urogenital system. Dev Cell. 2005;9(2):283–92.

4. Park JS, Valerius MT, and McMahon AP. Wnt/beta-catenin signaling regulates nephron induction during mouse kidney development. Development. 2007;134(13):2533–9.

5. Park JS, Ma W, O’Brien LL, Chung E, Guo JJ, Cheng JG, et al. Six2 and Wnt regulate self-renewal and commitment of nephron progenitors through shared gene regulatory networks. Dev Cell. 2012;23(3):637–51.

6. Karner CM, Das A, Ma Z, Self M, Chen C, Lum L, et al. Canonical Wnt9b signaling balances progenitor cell expansion and differentiation during kidney development. Development. 2011;138(7):1247–57.

7. Das A, Tanigawa S, Karner CM, Xin M, Lum L, Chen C, et al. Stromal-epithelial crosstalk regulates kidney progenitor cell differentiation. Nat Cell Biol. 2013;15(9):1035–44.

8. Kopan R, and Ilagan MX. The canonical Notch signaling pathway: unfolding the activation mechanism. Cell. 2009;137(2):216–33.

9. Chung E, Deacon P, Marable S, Shin J, and Park JS. Notch signaling promotes nephrogenesis by downregulating Six2. Development. 2016;143(21):3907–13.

10. Chung E, Deacon P, and Park JS. Notch is required for the formation of all nephron segments and primes nephron progenitors for differentiation. Development. 2017;144(24):4530–9.

11. Jeong HW, Jeon US, Koo BK, Kim WY, Im SK, Shin J, et al. Inactivation of Notch signaling in the renal collecting duct causes nephrogenic diabetes insipidus in mice. J Clin Invest. 2009;119(11):3290–300.

12. Mukherjee M, deRiso J, Otterpohl K, Ratnayake I, Kota D, Ahrenkiel P, et al. Endogenous Notch Signaling in Adult Kidneys Maintains Segment-Specific Epithelial Cell Types of the Distal Tubules and Collecting Ducts to Ensure Water Homeostasis. J Am Soc Nephrol. 2018.

13. Boyle SC, Liu Z, and Kopan R. Notch signaling is required for the formation of mesangial cells from a stromal mesenchyme precursor during kidney development. Development. 2014;141(2):346–54.

14. Lin EE, Sequeira-Lopez ML, and Gomez RA. RBP-J in FOXD1+ renal stromal progenitors is crucial for the proper development and assembly of the kidney vasculature and glomerular mesangial cells. Am J Physiol Renal Physiol. 2014;306(2):F249–58.

15. Park J, Shrestha R, Qiu C, Kondo A, Huang S, Werth M, et al. Single-cell transcriptomics of the mouse kidney reveals potential cellular targets of kidney disease. Science. 2018;360(6390):758–63.

16. Gribouval O, Gonzales M, Neuhaus T, Aziza J, Bieth E, Laurent N, et al. Mutations in genes in the renin-angiotensin system are associated with autosomal recessive renal tubular dysgenesis. Nat Genet. 2005;37(9):964–8.

17. Gribouval O, Moriniere V, Pawtowski A, Arrondel C, Sallinen SL, Saloranta C, et al. Spectrum of mutations in the renin-angiotensin system genes in autosomal recessive renal tubular dysgenesis. Hum Mutat. 2012;33(2):316–26.

18. Deacon P, Concodora CW, Chung E, and Park J-S. β-catenin regulates the formation of multiple nephron segments in the mouse kidney. Scientific Reports. 2019;9(1):15915.

19. Marable SS, Chung E, and Park JS. Hnf4a Is Required for the Development of Cdh6-Expressing Progenitors into Proximal Tubules in the Mouse Kidney. J Am Soc Nephrol. 2020.

20. Cho EA, Patterson LT, Brookhiser WT, Mah S, Kintner C, and Dressler GR. Differential expression and function of cadherin-6 during renal epithelium development. Development. 1998;125(5):803–12.

21. Marable SS, Chung E, Adam M, Potter SS, and Park JS. Hnf4a deletion in the mouse kidney phenocopies Fanconi renotubular syndrome. JCI Insight. 2018;3(14).

22. Soriano P. Abnormal kidney development and hematological disorders in PDGF beta-receptor mutant mice. Genes Dev. 1994;8(16):1888–96.

23. Grigorieva IV, Oszwald A, Grigorieva EF, Schachner H, Neudert B, Ostendorf T, et al. A Novel Role for GATA3 in Mesangial Cells in Glomerular Development and Injury. J Am Soc Nephrol. 2019;30(9):1641–58.

24. Martinez MF, Medrano S, Brown EA, Tufan T, Shang S, Bertoncello N, et al. Super-enhancers maintain renin-expressing cell identity and memory to preserve multi-system homeostasis. J Clin Invest. 2018;128(11):4787–803.

25. Yanai K, Saito T, Kakinuma Y, Kon Y, Hirota K, Taniguchi-Yanai K, et al. Renin-dependent cardiovascular functions and renin-independent blood-brain barrier functions revealed by renin-deficient mice. J Biol Chem. 2000;275(1):5–8.

26. Xu D, Borges GR, Grobe JL, Pelham CJ, Yang B, and Sigmund CD. Preservation of intracellular renin expression is insufficient to compensate for genetic loss of secreted renin. Hypertension. 2009;54(6):1240–7.

27. Takahashi N, Lopez ML, Cowhig JE, Jr., Taylor MA, Hatada T, Riggs E, et al. Ren1c homozygous null mice are hypotensive and polyuric, but heterozygotes are indistinguishable from wild-type. J Am Soc Nephrol. 2005;16(1):125–32.

28. Wu H, Uchimura K, Donnelly EL, Kirita Y, Morris SA, and Humphreys BD. Comparative Analysis and Refinement of Human PSC-Derived Kidney Organoid Differentiation with Single-Cell Transcriptomics. Cell Stem Cell. 2018;23(6):869–81 e8.

29. Takasato M, Er PX, Chiu HS, Maier B, Baillie GJ, Ferguson C, et al. Kidney organoids from human iPS cells contain multiple lineages and model human nephrogenesis. Nature. 2015.

30. Homan KA, Gupta N, Kroll KT, Kolesky DB, Skylar-Scott M, Miyoshi T, et al. Flow-enhanced vascularization and maturation of kidney organoids in vitro. Nat Methods. 2019;16(3):255–62.

31. Kobayashi A, Mugford JW, Krautzberger AM, Naiman N, Liao J, and McMahon AP. Identification of a multipotent self-renewing stromal progenitor population during mammalian kidney organogenesis. Stem cell reports. 2014;3(4):650–62.

32. Srinivas S, Watanabe T, Lin CS, William CM, Tanabe Y, Jessell TM, et al. Cre reporter strains produced by targeted insertion of EYFP and ECFP into the ROSA26 locus. BMC Dev Biol. 2001;1:4.

33. Madisen L, Zwingman TA, Sunkin SM, Oh SW, Zariwala HA, Gu H, et al. A robust and high-throughput Cre reporting and characterization system for the whole mouse brain. Nat Neurosci. 2010;13(1):133–40.

34. Yang X, Klein R, Tian X, Cheng HT, Kopan R, and Shen J. Notch activation induces apoptosis in neural progenitor cells through a p53-dependent pathway. Dev Biol. 2004;269(1):81–94.

35. McCright B, Lozier J, and Gridley T. Generation of new Notch2 mutant alleles. Genesis. 2006;44(1):29–33.

36. Tanigaki K, Han H, Yamamoto N, Tashiro K, Ikegawa M, Kuroda K, et al. Notch-RBP-J signaling is involved in cell fate determination of marginal zone B cells. Nat Immunol. 2002;3(5):443–50.

37. Hori K, Cholewa-Waclaw J, Nakada Y, Glasgow SM, Masui T, Henke RM, et al. A nonclassical bHLH Rbpj transcription factor complex is required for specification of GABAergic neurons independent of Notch signaling. Genes Dev. 2008;22(2):166–78.

38. Schmahl J, Rizzolo K, and Soriano P. The PDGF signaling pathway controls multiple steroid-producing lineages. Genes Dev. 2008;22(23):3255–67.

39. Amsen D, Antov A, Jankovic D, Sher A, Radtke F, Souabni A, et al. Direct regulation of Gata3 expression determines the T helper differentiation potential of Notch. Immunity. 2007;27(1):89–99.

40. Adam M, Potter AS, and Potter SS. Psychrophilic proteases dramatically reduce single-cell RNA-seq artifacts: a molecular atlas of kidney development. Development. 2017;144(19):3625–32.

41. Hao Y, Hao S, Andersen-Nissen E, Mauck WM, 3rd, Zheng S, Butler A, et al. Integrated analysis of multimodal single-cell data. Cell. 2021;184(13):3573–87 e29.

42. Hafemeister C, and Satija R. Normalization and variance stabilization of single-cell RNA-seq data using regularized negative binomial regression. Genome Biol. 2019;20(1):296.

