## Supplemental Figure 1 for "Interstitial Notch signaling regulates nephron development via the Gata3-Renin axis in the mouse kidney"

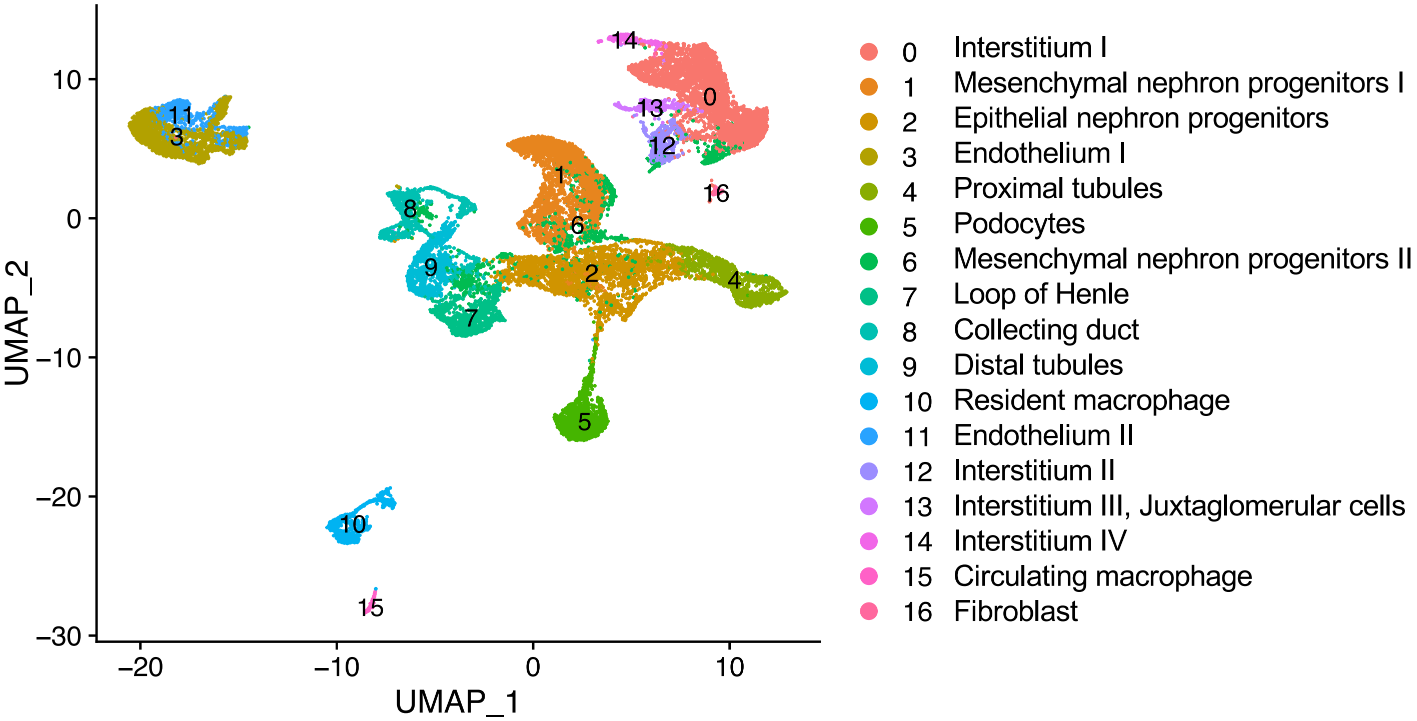


**Supplemental Figure 1.** scRNA-seq analysis of the interstitial *Rbpj* mutant and control kidneys at E18.5. Clustering of single cell expression profiles shows 17 cell types.
