## Supplemental Figure 2 for "Interstitial Notch signaling regulates nephron development via the Gata3-Renin axis in the mouse kidney"

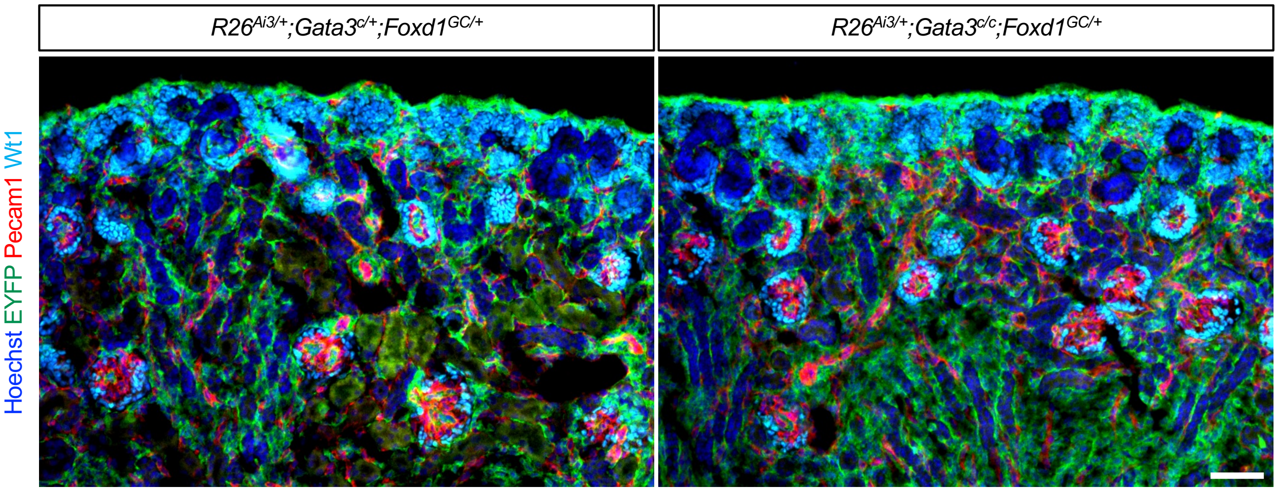


**Supplemental Figure 2.** EYFP$^{+}$ cells are present in most of the glomeruli in the *Gata3* mutant kidney by *Foxd1Cre*, suggesting that the mesangial cell defect is minimal. Stage P0; Scale bar 100 μm.
