## Supplemental Table 1 for "Interstitial Notch signaling regulates nephron development via the Gata3-Renin axis in the mouse kidney"

**Supplemental Table 1. Antibodies used for immunofluorescence**

Antigen	Vendors	Catalog #	Host	dilution

GFP	Aves Labs	GFP-1020	chick	1:500

Pecam1	Santa Cruz	sc-18916	rat IgG2a	1:500

Wt1	Santa Cruz	sc-7385	mouse	1:100

Wt1	Abcam	ab89901	rabbit	1:500

Cdh6	MilliporeSigma	HPA007047	rabbit	1:200

Hnf4a	Abcam	ab41898	mouse IgG2a	1:500

biotin-LTL	Vector Laboratories	B-1325	1:900

FITC-LTL	Vector Laboratories	FL-1321	1:500

Pdgfrb	Thermo Fisher	50-112-2656	rat IgG2a	1:500

Ren1	MilliporeSigma	HPA005131	rabbit	1:200
